# A novel non-invasive method to detect gut barrier related changes during a gastrointestinal nematode infection

**DOI:** 10.1101/496760

**Authors:** Norus Ahmed, Nicole Affinass, Emanuel Heitlinger, Anja A. Kühl, Natasa Xenophontos, Victor Hugo Jarquin, Jenny Jost, Svenja Steinfelder, Susanne Hartmann

## Abstract

Currently, methods for monitoring changes of gut barrier integrity and the associated immune response via non-invasive means are limited. Therefore, we aimed to develop a novel non-invasive technique to investigate immunological host responses representing gut barrier changes in response to infection. We identified the mucous layer on feces from mice to be mainly composed of exfoliated intestinal epithelial cells. Expression of RELM-β, a gene prominently expressed in intestinal nematode infections, was used as an indicator of intestinal cellular barrier changes to infection. RELM-β was detected as early as 6 days post-infection (dpi) in exfoliated epithelial cells. Interestingly, RELM-β expression also mirrored the quality of the immune response, with higher amounts being detectable in a secondary infection and in high dose nematode infection in laboratory mice. This technique was also applicable to captured worm-infected wild house mice. We have therefore developed a novel non-invasive method reflecting gut barrier changes associated with alterations in cellular responses to a gastrointestinal nematode infection.

## Introduction

Soil transmitted, intestinal nematodes affect around 24% of the world’s population (1) and are prevalent in wild animals. The majority of parasitic helminths live in the gut and are in close contact with the host’s epithelial cell (2), representing an important barrier during infection (3). The gut barrier is composed of specific enterocytes that secrete among others mucins the cysteine rich cytokine RELM-β and antimicrobial peptides, together forming the epithelial barrier during infection (4). This barrier defends against pathogen invasion but also leads to the activation of the mucosal immune system in the underlying lamina propria. In helminth infections the activation of the mucosal immune system leads to the activation of a T helper 2 (Th2) immune response. During a mouse gastrointestinal (GI) infection with *Heligmosomoides polygyrus*, the Th2 immune response is characterized by the expression of the transcription factor GATA-3, the cysteine-rich cytokine resistin-like molecule-beta (RELM-β), the cytokines interleukin (IL)-4, IL-5, IL-9, IL-13 and the antibodies IgE and IgG1 (5,6,7). This Th2 response leads to reduced worm fecundity and parasite expulsion (8,9). In this scenario, RELM-β has been shown to play multiple roles in different aspects of host defenses. It aids in spatial separation of the colonic epithelium and the microbiota by acting as a bactericidal protein (10,11). Additionally, RELM-β plays a role in immune regulation and host defenses against intestinal nematode infections. In a *H. polygyrus* infection it prevents worm feeding on host tissues and contributes to the weep and sweep response (12,13). Notably, during *H. polygyrus* infection significant changes in the composition of the gut microbiota have been documented (13,14).

Currently, different diseases are detectable using non-invasive techniques that utilize samples from urine, saliva, and stool. Urine samples can be used to detect selected viral, bacterial and parasitic diseases (15,16), including typhoid fever (17) and eggs of the blood fluke *Schistosoma haematobium* (18). Saliva has been a useful source in the early detection of foot and mouth disease in wild boar (19). Moreover, bacterial infections such as *Helicobacter pylori* (20) and the parasite *Plasmodium falciparum* are detected non-invasively using saliva from infected patients (16). Stool is regularly used to monitor numerous wildlife populations, including detection of virus infections in gorillas (21) or wild apes (22), bacterial shedding in the European badger as well as helminth eggs in Asian elephants (23,24) *or African buffalos* (25). In addition, stool is also used in humans to detect cytomegalovirus DNA instead of using mucosal biopsies (26).

In gastrointestinal helminth infections various helminth eggs are detectable using stool, however this method does not detect the early stages of infection. Neither does it investigate host parameters associated with disease, such as gut barrier related changes. Therefore, a non-invasive sampling technique that provides further information about cellular changes during infection and enables the monitoring of disease progression is urgently required for both laboratory and field settings. A non-invasive assessment reflecting cellular and immunological gut parameters would provide a better understanding of pathogen burdens, detection of communities prone to diseases and the identification of immunologically naïve populations (15,27).

Here, we describe a novel non-invasive method that uses exfoliated intestinal cells to monitor cellular and immunological changes of the gut barrier in response to infection. We establish and demonstrate this method in laboratory mice infected with the small intestinal nematode *H. polygyrus*. By comparing acute versus chronic infection, dose-dependent responses and reinfection after abrogation of infection, we illustrate the potential of detecting cellular responses after gut barrier changes using RNA extracted from exfoliated intestinal cells. In addition, we applied this method to wild mouse stool samples. Thus, this study uses the gene expression from exfoliated cells present on stool to detect changes in gut barrier function due to infectious diseases by non-invasive means.

## Materials and Methods

### Animals

Female BALB/c mice were used (8 weeks old; purchased from Janvier, Saint Berthevin, France). The experiments were performed followed the National Animal Protection Guidelines and were approved by the German Animal Ethics Committee for the protection of animals (H0438/17, G0113/15, G0176/16).

### Infection

*H. polygyrus* was maintained by serial passage in C57BL/6 mice, described previously (Rausch et al., 2008). Mice aged 8 weeks were infected by oral gavage with either 20 or 200 infective L3 larvae diluted in drinking water. On either 14 or 35 days post infection (dpi) mice were sacrificed by isofluorane inhalation. Mice were treated during the acute phase of infection (day 14 and day 15) with 2 mg/animal pyrantel pamoate for adult worm expulsion (Sigma, St. Louis, MO, USA) in 200 μL water.

### Fecal collection and storage

4-10 freshly excreted fecal pellets were collected from mice. Fecal pellets were collected in cryotubes, placed into liquid nitrogen and stored at −80°C until processing.

### Wild mice sampling

House mice (*Mus musculus*) were captured in autumn 2017 using live traps (approval number 35 – 2014–2347). Traps were set overnight in farms and private properties in the state of Brandenburg in Germany. All animals were transferred to individual cages and remained there until fecal samples were collected on the following day. Fresh fecal samples were kept in liquid nitrogen during transportation and maintained at −80°C until processing. Mice were euthanized, digestive tracts were dissected and helminths were identified and counted under a binocular microscope.

### Intestinal exfoliated epithelial cell extraction

Flotation method: Falcon tubes (15 mL) were filled with 3 mL PBS and put on ice. Faecal pellets previously stored at −80°C were used and one pellet was placed individually into each falcon tube. Tubes were placed on a rocker for 45 minutes at 4°C. Tubes were then turned upright on ice for a further 20-30 minutes to allow cells to loosen up. Another set of falcon tubes were prefilled with 3 mL PBS and stored on ice. Individual falcon tubes containing a single fecal pellet were slowly inverted until the layer of epithelial cells started to detach from the fecal pellet. The inverting force on the falcon tubes was slowly increased until the layer floated off. Epithelial cells could also be removed using a pipette while rotating the fecal pellet to completely peel off the layer. These cells were then collected using a pipette and placed into the freshly prepared falcon tubes to remove any fecal debris. This material was subsequently used for RNA extraction (Analytik Jena, Jena, Germany).

Alternative field method: Fecal pellets were placed in 2 mL tubes filled with 1 mL RNA later (Sigma-Aldrich, St. Louis, MO, USA) and stored at −20°C. For the removal of exfoliated epithelial cells samples were placed in 3 mL PBS filled falcon tubes and left upright for 60-90 minutes before inverted until cells float off. A rocker was not required for this method.

### Histopathology

Mucus preparations taken from feces were fixed in formalin and embedded in paraffin. Paraffin sections of 1-2 μm thickness were cut, dewaxed and stained histochemically with hematoxylin and eosin (H&E) for overview and with periodic acid-Schiff (PAS) for mucus. Sections were cover slipped with corbit balsam (Hecht, Germany). For immunohistochemical detection of epithelial cells, paraffin sections were dewaxed and subjected to a heat-induced epitope retrieval step prior to incubation with anti-EpCAM (clone E6V8Y, Cell Signaling). For detection, EnVision+ System-HRP Labelled Polymer Anti-Rabbit (Agilent Technologies) was employed. Nuclei were counterstained with hematoxylin (Merck). Negative controls were performed by omitting the primary antibody. Images were acquired using the AxioImager Z1 microscope (Carl Zeiss MicroImaging, Inc.). All evaluations were performed in a blinded manner.

### Realtime PCR

RNA was isolated from exfoliated cells, intestinal tissue sections and whole fecal pellets that were previously stored at −80 °C via homogenization in RNA lysis buffer. The homogenized exfoliated cells, fecal pellets and tissue samples were centrifuged and supernatants were treated with a innuPREP RNA kit (Analytik Jena, Jena, Germany) following manufacturer’s instructions. 2 μg of RNA was reverse transcribed to cDNA using a High Capacity RNA to cDNA kit (Applied Biosystems, Foster City, CA). The relative expression of β-actin, resistin-like molecule-beta (RELM-β), IL-25, IL-33 and TSLP was determined by Real Time PCR using 10 ng of cDNA and the FastStart Universal SYBR Green Master Mix (Roche, Basel, Switzerland). Relative gene expression of two-three technical replicates is shown as mean. Primer pairs used for gene amplification were as follows: β-actin (*Actb*) forward: GGCTGTATTCCCCTCCATCG, reverse: CCAGTTGGTAACAATGCCATGT, Relm-β (*Retnlb*) forward: GGCTGTGGATCGTGGGATAT, reverse: GAGGCCCAGTCCATGACTGA, IL-33 (*Il33*) forward: AGGAAGAGATCCTTGCTTGGCAGT, reverse: ACCATCAGCTTCTTCCCATCCACA, IL-25 (*Il25*) forward: ACAGGGACTTGAATCGGGTC, reverse: TGGTAAAGTGGGACGGAGTTG, TSLP (*Tslp*) forward: T GGT AAAGT GGGACGGAGT T G and reverse: TGTGCCATTTCCTGAGTACCGTCA. GAPDH (*Gapdh*) forward: TGGATTTGGACGCATTGGTC, reverse: TTTGCACTGGTACGTGTTGAT; HPRT (*Hprt*) forward: TCCTCCTCAGACCGCTTTT, reverse: TTTCCAAATCCTCGGCATAATGA; GUS-β (*Gusb*) forward: GCTCGGGGCAAATTCCTTTC, reverse: CTGAGGTAGCACAATGCCCA. The mRNA expression was normalized to the β-actin using the Δ-ct value and calculated by Roche Light Cycler 480 software.

### Fecal egg counts

Fecal pellets were collected and placed in glass tubes. Fecal pellets were homogenized and mixed with 1 mL tap water. Next, 6 mL of a saturated salt solution (NaCl) was added to the homogenized sample. The McMaster chamber was then filled with sample and eggs were counted.

### Flow Cytometry

For surface and intracellular staining, the following monoclonal antibodies were used: CD4 (PerCP) (RM4-5) from BioLegend (Biozol); IL-13 (Alexa 488) (eBio13A); Foxp3 Alexa 488 (FJK-16s); GATA-3 (eFluor 660) (TWAJ) and Dead Cell Exclusion Marker (DCE) (efluor 780) all from eBioscience, San Diego, CA, USA.

### Statistics

Experiments are displayed as either mean ± SD or mean ± SEM as indicated. Statistical analysis was completed using GraphPad Prism software (La Jolla, CA, USA). Significance was determined as indicated using the Kruskal-Wallis with Dunn’s multiple comparison test or the Mann Whitney U Test. Also, multiple t tests were corrected for multiple comparisons using the Holm-Sidak method. Significance was measured as \**P* ≤ 0.05, \*\**P* ≤ 0.01, and \*\*\**P* ≤ 0.001.

## Results

### Identification of epithelial cells and mucus in a layer of exfoliated intestinal cells from stool

A layer of exfoliated cells from stool (Figure 1A) was analyzed using different histological techniques. A thick mucus layer is apparent in an H&E stain (1B) and in a PAS-staining (1C) with cells either attached to the mucus or in between mucinous strands. These cells displayed a broad cytoplasm suggesting that they are of epithelial origin and were probably scaled off during defecation. Immunohistochemical detection of EpCAM expression confirmed that the majority of these cells are epithelial cells (Figure 1D and E). Thus, epithelial cells can be retrieved from the surface of stool (Figure 1F).

**Figure 1.**
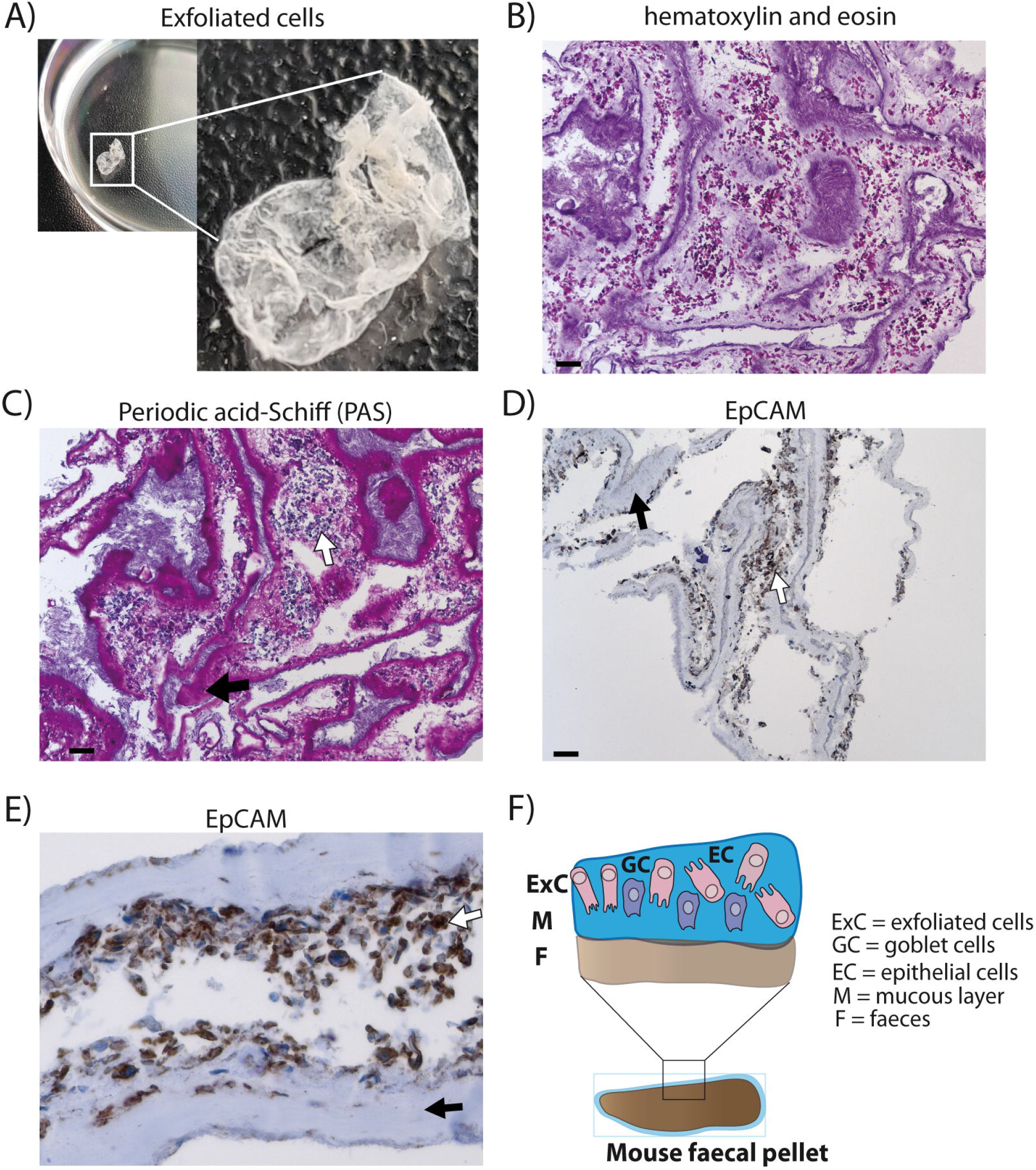
Analysis of exfoliated intestinal epithelial cells isolated from murine stool. A) A layer of exfoliated cells removed from the surface of mouse stool in PBS. B) Hematoxylin and eosin (H&E) stain of exfoliated cells extracted from the surface of stool at 100x magnification (scale bar 100μm). C) Periodic acid-Schiff (PAS) staining for mucus detection in exfoliated cells at 100x magnification (scale bar 100μm). D) EpCAM staining identifying epithelial cells using exfoliated cells (brown staining) at 100x magnification (scale bar 100μm) E) EpCAM staining of epithelial cells at 400x magnification. F) Schematic diagram of stool surface with exfoliated cells. C-E) White arrows display cells and black arrows highlight mucus.

### Quantification of RNA isolated from exfoliated intestinal cells

Currently there is a lack of information on cellular and immunological changes of the gut barrier after infection using non-invasive techniques. Here, we established a method allowing the quantification of RNA from exfoliated intestinal cells. Exfoliated cells showed a lower Cycle threshold (Ct) value for the amplification of RELM-β compared to homogenized whole fecal pellets. The Ct value is defined as the required number of cycles for the signal to exceed the fluorescence threshold in quantitative PCR (Supplementary table 1). Furthermore, exfoliated cells showed very low Ct value variation between technical triplicates, whereas homogenized fecal pellets sometimes displayed a difference of 10 in Ct values within triplicates. We attributed the lower quality of the homogenized samples to higher degradation of mRNA based on the differences in Ct values within triplicates. Therefore, we regard the extraction of exfoliated cells to be superior to whole fecal pellet homogenization. To identify the ideal housekeeping gene, the different housekeeping genes β-actin, GAPDH, HPRT and GUS-β were analyzed. β-actin, GAPDH, GUS-β and HPRT were tested using 10 ng cDNA and 100 ng cDNA (Supplementary table 2). While expression levels of all four housekeeping genes correlated with each other, β-actin produced a lower Ct value at 10 ng compared to the other three housekeeping genes at 100 ng. Thus, β-actin was the housekeeping gene of choice throughout. β-actin was also detectable using fecal samples stored at −20°C in RNA later for 30 days (data not shown).

### RELM-β expression in exfoliated cells during acute and chronic intestinal helminth infection

Depending on the infection, the immune responses vary in terms of quantity and quality. Here, we aimed to quantify mRNA expression of a selection of immune genes typical for a Th2 type response against helminth infections. We aimed to obtain infection-specific data using exfoliated intestinal cells isolated from stool. To establish this technique, a murine infection model with the small intestinal nematode *H. polygyrus* was used. During the early acute phase of *H. polygyrus* infection the larvae enter the intestinal tissue and develop into the fourth larval (L4) stage, then re-enter the lumen of the gut. In response to the tissue invasive phase and resulting tissue damage the epithelium-derived cytokines IL-25, IL-33 and TSLP are released (28). During an acute *H. polygyrus* infection (14 dpi), our data shows that mRNA expression of these tissue-derived cytokines from exfoliated cells were insufficiently detected and showed high variability between triplicates (Figure S1A-C). We thus investigated whether exfoliated cells could be used to detect the Th2 cysteine-rich cytokine RELM-β. RELM-β is produced by intestinal goblet cells that significantly increase during intestinal helminth infection (29). Interestingly, RELM-β expression was significantly detected as early as day 6 post *H. polygyrus* infection (Figure 2A). This increase of expression was also significant at day 8 and 10, steadily decreasing thereafter, making it detectable throughout the course of an acute infection. In the case of chronic helminth infection, RELM-β expression continued to decrease from day 21 to levels similar to day 0 (Figure 2B). As a method to confirm infection, helminth eggs were counted in parallel during the acute and chronic phase (Figure 2C and D). As expected, egg counts and RELM-β mRNA expression peaked at different time points but overall mirrored a similar course of infection. Of note, the RELM-β detection in exfoliated cells directly correlated with the local gut Th2 immune response investigated via flow cytometry during the course of infection. An increase in the Th2 transcription factor GATA-3+ in CD4^+^ T cells in mesenteric lymph nodes (mLN) peaked at day 8 post *H. polygyrus* infection (Figure 2E). The same was observed for the Th2 cytokine IL-13 (Figure 2F). When investigating the systemic immune response, we observed a delayed peak in the expression of GATA-3^+^ in CD4^+^ T cells at day 14 in the spleen, decreasing onwards to day 35 (Figure 2G). The same was observed for the IL-13 expression in the spleen indicating a delayed systemic response to infection (Figure 2H). Thus, the local immune response in the infection draining mLN mirrored RELM-β detection in exfoliated cells.

**Figure 2.**
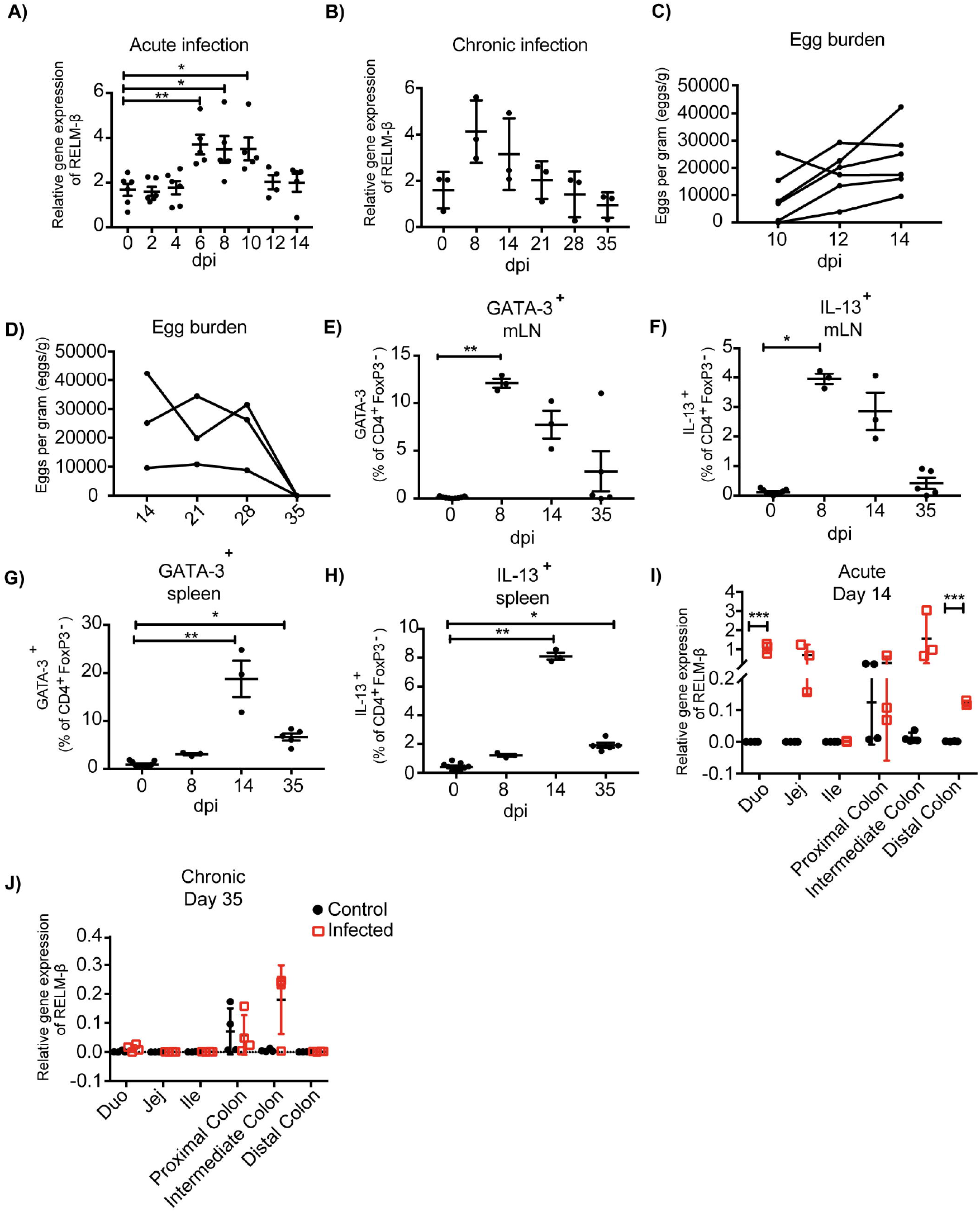
Immunological and parasitological analysis of murine stool after infection with *Heligmosomoides polygyrus*. BALB/c mice were infected orally with 200 infectious L3 stage larvae of *H. polygyrus*. Parameters were measured at different time points and in different regions of the intestine. A) Relative gene expression of RELM-β in exfoliated intestinal cells during acute infection (day 0 - 14 dpi). B) RELM-β gene expression in exfoliated cells until chronicity of infection (day 0 – 35 dpi), shown as mean ± SD. C) Fecal egg counts during acute *H. polygyrus* infection (day 14 dpi). D) Fecal egg counts during chronic *H. polygyrus* infection (day 14 – 35 dpi). Frequency of CD4+ T cells expressing GATA-3 (E) and IL-13 (F) in mesenteric lymph nodes (mLN). G) Frequencies of CD4+ T cell expressing GATA-3 and IL13 (H) in spleen. I) and J), RELM-β expression in intestinal tissue (duodenum, jejunum, ileum, proximal colon, intermediate colon and distal colon) at 14 dpi and 35 dpi, respectively. All relative gene expression analysis is compared to β-actin using 10ng cDNA. Data from panel (A) is pooled from two independent experiments with n=4-10 fecal pellets from 4-6 animals. Panels (B) is representative of two independent experiments, n=3. Panel (C) is pooled from two independent experiments, n=6. Panel (D-H) is representative of two independent experiments, n=3. Panel (I and J) is representative of two independent experiments, n=3-4. Panel A shown as mean ± SEM, \**P* ≤ 0.05, \*\**P* ≤ 0.01, and \*\*\**P* ≤ 0.001. Panel B-H shown as mean ± SD; A-H) Statistical analysis was performed using the Kruskal-Wallis with Dunn’s multiple comparison test, I-J) multiple t tests and corrected for multiple comparisons using the Holm-Sidak method, \**P* ≤ 0.05, \*\**P* ≤ 0.01, and \*\*\**P* ≤ 0.001. dpi: days post infection

To verify the observed mRNA levels in exfoliated cells, we examined the cytokine expression in host tissue along the gut. In parallel, intestinal tissues from the duodenum, jejunum, ileum, proximal colon, intermediate colon and distal colon were analyzed to decipher the expression of the tissue derived cytokines IL-25, IL-33, TSLP (S1D-F) and the goblet cell produced RELM-β. Interestingly, at 14 days post-infection a significant RELM-β signal was detectable in the duodenum and distal colon during acute *H. polygyrus* infection (Figure 2I). However, RELM-β mRNA was not observed in intestinal tissue at day 35 post infection (Figure 2J) correlating with the RELM-β detection in exfoliated cells. This data indicates a reduced Th2 immune response against the worm infection, probably due to advanced clearance of the parasite.

In summary, RELM-β mRNA as an indicator for gut barrier changes was detectable as early as day 6 post nematode infection in exfoliated intestinal cells. The non-invasively detected RELM-β expression mirrored the immune responses at the gut barrier. Thus, exfoliated epithelial cells from stool can be used to detect infection.

### Pathogen dose-dependent expression of the cysteine-rich cytokine RELM-β in exfoliated epithelial cells

Next, we aimed to investigate if the detection of the cysteine-rich cytokine RELM-β correlated to the intensity of infection. Changes in the gene expression of RELM-β in exfoliated gut cells were studied using different infection dosages. Mice were infected with either a low dose infection with 20 L3 of *H. polygyrus* or a high dose infection with 200 L3 and adult worm burden was assessed at day 14 (Figure 3A). The high dose infection was reflected by a higher RELM-β expression compared to a low dose infection. The expression of RELM-β during the 200 L3 infection was significantly higher compared to a low dose infection at day 8 p.i. (Figure 3B). This effect was most prominent at day 8 but a trend was detected from day 4 p.i. onwards. Thus, RELM-β expression in exfoliated cells does mimic the intensity of an intestinal infection.

**Figure 3.**
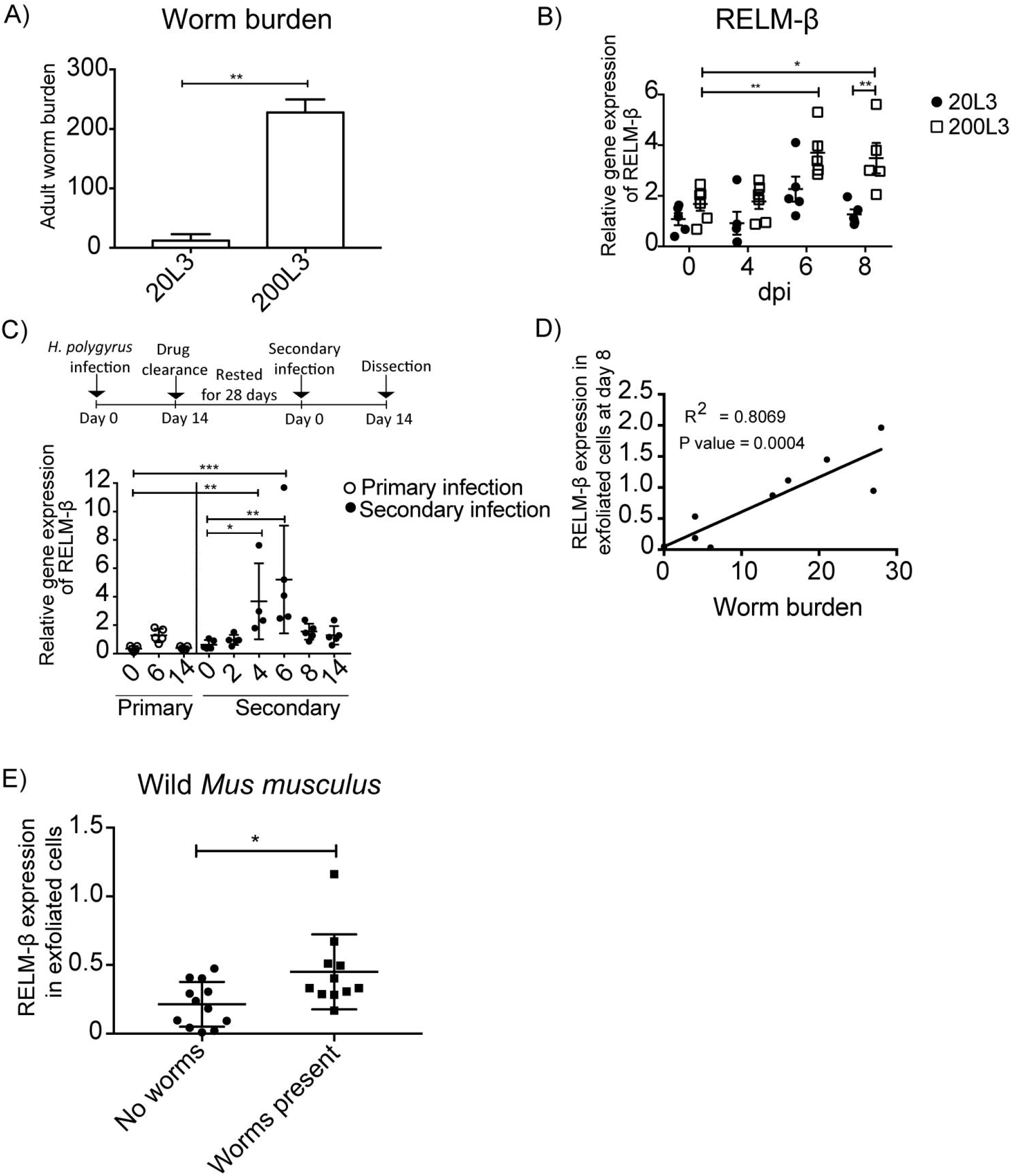
RELM-β expression in exfoliated cells reflects the intensity and type of infection. BALB/c mice were infected orally with either a low dose (20 L3) or a high dose (200 L3) *H. polygyrus* larvae. A) Worm burden assessed at 14 dpi. B) RELM-β relative gene expression during early *H. polygyrus* infection until day 8 dpi after low dose (20 L3) and high dose (200 L3) infection. C) BALB/c mice were infected orally with 200 L3 *H. polygyrus*, treated with pyrantel pamoate for worm clearance and 28 days later reinfected with 200 L3. RELM-β gene expression shown during primary and secondary infection. D) Linear regression comparing RELM-β gene expression at day 8 in exfoliated cells with low worm burden. Panels (A) is representative from two independent experiments with n = 3-5 animals and shows the mean ± SD. B) is pooled from two independent experiments with n = 4-6 animals, data shown as mean ± SEM. All relative gene expression analysis is compared to β-actin using 10ng cDNA. Panel (C) is representative of two independent experiments, n=4-5. Panel (D) is pooled from two independent experiments, n=10. Statistical analysis using A) Mann Whitney U Test, B) multiple t tests and corrected for multiple comparisons using the Holm-Sidak method, C) was performed using the Kruskal-Wallis with Dunn’s multiple comparison test. E) Mann Whitney U Test, \**P* ≤ 0.05, \*\**P* ≤ 0.01, and \*\*\**P* ≤ 0.001. All relative gene expression analysis is compared to β-actin.

In nature, animals frequently get re-infected with helminths. Therefore, we asked if our new non-invasive method is suitable to detect the expression of RELM-β as a marker for gut barrier changes after drug clearance of *H. polygyrus* and reinfection. Mice were re-infected 28 days after drug clearance with 200 L3 infective larvae. RELM-β mRNA was significantly detectable in exfoliated gut cells at day 4 and day 6 in mice re-infected with *H. polygyrus* (Figure 3C). Interestingly, a significantly positive linear relationship was observed between the expression of RELM-β in exfoliated cells and adult worm counts (Figure 3D).

### Analysis of intestinal exfoliated cell gene expression in wild mice samples

In order to analyze the potential of the non-invasive cytokine detection method with wildlife animals, we tested stool samples collected from wild *Mus musculus* mice captured in Brandenburg, Germany. RELM-β gene expression was analyzed in exfoliated intestinal cells isolated from fecal pellets from wild mice with no worms detectable in the gut. The data was then compared to data obtained from wild mice with different helminths present at the time of collection and dissection (Supplementary Table 3). We also observed a significant increase of RELM-β expression in samples from wild mice with worms (Figure 3E). Our study with samples from wild animals shows that RELM-β is detectable during an active infection with intestinal helminths.

## Discussion

Here we investigated whether exfoliated intestinal cells released with feces mirror immunological host responses corresponding to gut barrier changes during a gastrointestinal nematode infection with *H. polygyrus*. Investigation into infection related genes revealed the goblet cell-secreted RELM-β as a reliable marker reporting gut barrier changes due to acute worm infection. RELM-β was significantly detectable during the early stages of an acute *H. polygyrus* infection and decreased steadily thereafter. Intriguingly the mRNA levels of RELM-β in exfoliated cells reflected the local GATA-3 and IL-13 expression in mLN during infection. Our data suggests that detection of RELM-b in exfoliated gut cells not only detects gut barrier related changes but also reflects the local Th2 immune response.

In the intestine, epithelial cells represent an essential barrier that undergoes continuous homeostatic turnover replacing old cells and lost or damaged epithelial cells during infection (30,31). Additionally, epithelial cells are part of the mucosal immune response and aid in eliciting the prominent Th2 immune response during gastrointestinal helminth infection. Exfoliated cells are shed daily in feces and have been previously described to be of colonic origin. Interestingly during *H. polygyrus* infection goblet cells (GCs) play an important role in the secretion of RELM-β. The secretion of this cysteine-rich cytokine in intestinal tissue is essential for normal spontaneous worm expulsion (12). RELM-β acts on *H. polygyrus* by impairing feeding thus reducing protein and ATP content, leading to pathogen expulsion (29). Goblet cells also secrete mucin forming a mucosal surface barrier limiting interactions with luminal microbes (32). Additionally, during homeostasis GCs are highly abundant in the colon compared to the upper intestinal tract (13).

Importantly, our method is applicable in the field where cooling and proper storage of biological material is limited. We tested fecal pellets from wild mice freshly placed in RNA later™ and then stored at −20°C for 30 days. Interestingly, β-actin expression was detectable with a Ct value of 25, similar to flash frozen and −80°C stored fecal pellets (data not shown). Thus, this non-invasive method allows stool to be stored for longer periods of time before analysis, whereas egg counts need to be processed as soon as possible for a reliable representation of infection (33). This suggests that both egg counts and exfoliated cells should be used together to complement the information not only on the infection status but in addition the cellular host responses at the gut barrier.

Nematodes dwelling in the small intestine such as *H. polygyrus* have been shown to affect colonic permeability (34). This cellular response in the colon to an infection dwelling in the small intestine enables the detection of cellular responses using exfoliated cells likely originating from the colon. In addition, gastrointestinal nematodes have been described to alter the gut bacterial environment (13,35). Increase in abundance of gram-negative bacteria during *H. polygyrus* infection (13,14), accompanied by colonic barrier permeability (34) correlate with the increased early detection of RELM-β during infection in our study. Here, it is interesting to note that RELM-β is highly expressed (32) and plays multiple roles such as recruiting CD4^+^ T cells in *Citrobacter rodentium* infection (10). Additionally, RELM-β has been described as a bactericidal protein that kills gram-negative bacteria, limiting the association of bacteria with colonic tissues (11). Thus, the ability to detect RELM-β in exfoliated cells can possibly be attributed to the highly abundant goblet cells in the large intestine secreting RELM-β, to deal with gut barrier changes and the increase in bacteria during nematode infection. Thus, the previously described altered colonic permeability (34) and the increased gram-negative bacteria (13,14) explain the high expression of RELM-β and the detectability in exfoliated cells compared to the tissue cytokines.

Comparison of the tissue cytokines in different tissue compartments at 14 dpi confirmed no significant changes in IL-25, IL-33 and TSLP in the distal colon (Figure S1A-C). Additionally, expression of these tissue cytokines could not be detected using exfoliated cells from stool. The observed systemic tissue RELM-β expression and high expression in exfoliated cells displays a reliable marker that mirrors the local *H. polygyrus* Th2 immune responses non-invasively.

Additionally, this method can be utilized for other colonic infections and might display RELM-β as a marker of gut barrier changes.

In order to identify if exfoliated cells can be used as a measure of infection load, we tested a low versus high dose infection and identified significant differences in RELM-β expression. This method detected a difference in infection dose highlighting its potential as a marker of infection load and also reported the quality of the immune response in a secondary infection. A linear regression comparison of exfoliated cells compared to worm burden revealed a significantly positive relationship. Interestingly, RELM-β expression in exfoliated cells mirrored the kinetics of the local immune response in our study. Consequently, detection of RELM-β in exfoliated cells might be useful not only as an additional measure to egg counts but also as a means to monitor cellular gut barrier changes correlating to infection loads.

In wildlife, animals are exposed to a variety of pathogens and monitoring of the health status is urgently needed for wildlife conservation, zoonotic diseases and disease control. However, the only methods applicable to wildlife studies are non-invasive methods. Over time there have been many advances in different molecular biology techniques allowing for in depth research into different diseases. However, not much is known regarding cellular changes during infection in natural animal populations. We demonstrate here that our method using exfoliated cells to investigate infection related gut barrier changes works in both laboratory and wild animals.

In conclusion, we describe a novel method to measure cellular parameters using exfoliated cells from stool to detect infection-related gut barrier changes in samples from wildlife and experimental laboratory conditions. Further investigation into pathogen-specific infection markers is required. Thus far, our technique represents a potential addition to lab and field non-invasive techniques reflecting gut barrier changes. RELM-β detection in exfoliated cells can potentially provide a suitable marker for colonic inflammation in a variety of different diseases and colitis infections.

## Supporting information

## Acknowledgments

The authors thank Yvonne Weber, Marion Müller, Bettina Sonnenburg, Christiane Palissa, and Beate Anders for providing their excellent technical support and Ankur Midha for proofreading the article.

## Author contributions

N. Ahmed and NX performed all the exfoliated cell extractions and gene expression experiments. SH and SS conceptualized and designed the research. N. Affinass performed all the flow cytometry experiments and analysis. EH provided input into all wild mouse experimental data and analysis. AK performed all the histopathology experiments. VHJ was involved in the capturing and sample collection from wild mice and JJ was involved in the identification of worms in wild mouse samples. N. Ahmed and SH wrote the manuscript. All authors approved the final version of the manuscript.

---

**Supplementary figure 1. Relative gene expression of tissue cytokines in exfoliated cells and tissues following an *H. polygyrus* infection with 200 infectious L3 larvae.** Relative gene expression of A) IL-25, B) IL-33, C) TSLP in duodenum, jejunum, ileum, proximal colon, intermediate colon and distal colon at 14 dpi (acute infection). D) IL25, E) IL33, F) in exfoliated cells. G) IL-25, H) IL-33 I) TSLP in duodenum, jejunum, ileum, proximal colon, intermediate colon and distal colon at 35 dpi (chronic infection). All panels are representative of two independent experiments, n=3-5. Data is shown as mean ± SD. Statistical analysis was performed using the Kruskal-Wallis with Dunn’s multiple comparison test, \**P* ≤ 0.05, \*\**P* ≤ 0.01, and \*\*\**P* ≤ 0.001

**Supplementary table 1.** Real Time PCR cycle threshold (Ct) values of β-actin in exfoliated cells and homogenized stool using a 10 ng and 100 ng cDNA concentrations.

**Supplementary table 2.** Exfoliated cells were used to detect the Ct values of the different house keeping genes β-actin, GAPDH, HPRT and GUS-β using a 10 ng and 100 ng cDNA concentration

**Supplementary table 3.** Represents the location, sex, and infection status of wild *Mus musculus* mice captured in Brandenburg, Germany.

## References

1. Soil-transmitted helminth infections. Available at: https://www.who.int/news-room/fact-sheets/detail/soil-transmitted-helminth-infections [Accessed December 13, 2018]

2. McKay DM, Shute A, Lopes F. Helminths and intestinal barrier function. Tissue Barriers (2017) 5: doi:10.1080/21688370.2017.1283385

3. Linden SK, Sutton P, Karlsson NG, Korolik V, McGuckin MA. Mucins in the mucosal barrier to infection. Mucosal Immunol (2008) 1:183–197. doi:10.1038/mi.2008.5

4. McGuckin MA, Lindén SK, Sutton P, Florin TH. Mucin dynamics and enteric pathogens. Nat Rev Microbiol (2011) 9:265–278. doi:10.1038/nrmicro2538

5. Finkelman FD, Katona IM, Urban JF, Holmes J, Ohara J, Tung AS, Sample JV, Paul WE. IL-4 is required to generate and sustain in vivo IgE responses. J Immunol (1988) 141:2335–2341.

6. Urban JF, Katona IM, Finkelman FD. Heligmosomoides polygyrus: CD4+ but not CD8+ T cells regulate the IgE response and protective immunity in mice. Exp Parasitol (1991) 73:500–511.

7. Svetić A, Madden KB, Zhou XD, Lu P, Katona IM, Finkelman FD, Urban JF, Gause WC. A primary intestinal helminthic infection rapidly induces a gut-associated elevation of Th2-associated cytokines and IL-3. J Immunol (1993) 150:3434–3441.

8. Artis D, Wang ML, Keilbaugh SA, He W, Brenes M, Swain GP, Knight PA, Donaldson DD, Lazar MA, Miller HRP, et al. RELMbeta/FIZZ2 is a goblet cell-specific immune-effector molecule in the gastrointestinal tract. Proc Natl Acad Sci USA (2004) 101:13596–13600. doi:10.1073/pnas.0404034101

9. Owyang AM, Zaph C, Wilson EH, Guild KJ, McClanahan T, Miller HRP, Cua DJ, Goldschmidt M, Hunter CA, Kastelein RA, et al. Interleukin 25 regulates type 2 cytokine-dependent immunity and limits chronic inflammation in the gastrointestinal tract. J Exp Med (2006) 203:843–849. doi:10.1084/jem.20051496

10. Bergstrom KSB, Morampudi V, Chan JM, Bhinder G, Lau J, Yang H, Ma C, Huang T, Ryz N, Sham HP, et al. Goblet Cell Derived RELM-β Recruits CD4+ T Cells during Infectious Colitis to Promote Protective Intestinal Epithelial Cell Proliferation. PLoS Pathog (2015) 11:e1005108. doi:10.1371/journal.ppat.1005108

11. Propheter DC, Chara AL, Harris TA, Ruhn KA, Hooper LV. Resistin-like molecule β is a bactericidal protein that promotes spatial segregation of the microbiota and the colonic epithelium. Proc Natl Acad Sci USA (2017) 114:11027–11033. doi:10.1073/pnas.1711395114

12. Herbert DR, Yang J-Q, Hogan SP, Groschwitz K, Khodoun M, Munitz A, Orekov T, Perkins C, Wang Q, Brombacher F, et al. Intestinal epithelial cell secretion of RELM-beta protects against gastrointestinal worm infection. J Exp Med (2009) 206:2947–2957. doi:10.1084/jem.20091268

13. Rausch S, Held J, Fischer A, Heimesaat MM, Kühl AA, Bereswill S, Hartmann S. Small intestinal nematode infection of mice is associated with increased enterobacterial loads alongside the intestinal tract. PLoS ONE (2013) 8:e74026. doi:10.1371/journal.pone.0074026

14. Su C, Su L, Li Y, Chang J, Zhang W, Walker WA, Xavier RJ, Cherayil BJ, Shi HN. Helminth-Induced Alterations Of The Gut Microbiota Exacerbate Bacterial Colitis. Mucosal Immunol (2018) 11:144–157. doi:10.1038/mi.2017.20

15. Leendertz FH, Pauli G, Maetz-Rensing K, Boardman W, Nunn C, Ellerbrok H, Jensen SA, Junglen S, Christophe B. Pathogens as drivers of population declines: The importance of systematic monitoring in great apes and other threatened mammals. Biological Conservation (2006) 131:325–337. doi:10.1016/j.biocon.2006.05.002

16. Mfuh KO, Tassi Yunga S, Esemu LF, Bekindaka ON, Yonga J, Djontu JC, Mbakop CD, Taylor DW, Nerurkar VR, Leke RGF. Detection of Plasmodium falciparum DNA in saliva samples stored at room temperature: potential for a non-invasive saliva-based diagnostic test for malaria. Malar J (2017) 16:434. doi:10.1186/s12936-017-2084-5

17. Chaicumpa W, Ruangkunaporn Y, Burr D, Chongsa-Nguan M, Echeverria P. Diagnosis of typhoid fever by detection of Salmonella typhi antigen in urine. J Clin Microbiol (1992) 30:2513–2515.

18. Ibironke OA, Phillips AE, Garba A, Lamine SM, Shiff C. Diagnosis of Schistosoma haematobium by detection of specific DNA fragments from filtered urine samples. Am J Trop Med Hyg (2011) 84:998–1001. doi:10.4269/ajtmh.2011.10-0691

19. Mouchantat S, Haas B, Böhle W, Globig A, Lange E, Mettenleiter TC, Depner K. Proof of principle: non-invasive sampling for early detection of foot-and-mouth disease virus infection in wild boar using a rope-in-a-bait sampling technique. Vet Microbiol (2014) 172:329–333. doi:10.1016/j.vetmic.2014.05.021

20. Yu M, Zhang X-Y, Yu Q. Detection of oral Helicobacter Pylori infection using saliva test cassette. Pak J Med Sci (2015) 31:1192–1196. doi:10.12669/pjms.315.7626

21. D’arc M, Ayouba A, Esteban A, Learn GH, Boué V, Liegeois F, Etienne L, Tagg N, Leendertz FH, Boesch C, et al. Origin of the HIV-1 group O epidemic in western lowland gorillas. Proc Natl Acad Sci USA (2015) 112:E1343–1352. doi:10.1073/pnas.1502022112

22. Reed PE, Mulangu S, Cameron KN, Ondzie AU, Joly D, Bermejo M, Rouquet P, Fabozzi G, Bailey M, Shen Z, et al. A new approach for monitoring ebolavirus in wild great apes. PLoS Negl Trop Dis (2014) 8:e3143. doi:10.1371/journal.pntd.0003143

23. King HC, Murphy A, James P, Travis E, Porter D, Sawyer J, Cork J, Delahay RJ, Gaze W, Courtenay O, et al. Performance of a Noninvasive Test for Detecting Mycobacterium bovis Shedding in European Badger (Meles meles) Populations. J Clin Microbiol (2015) 53:2316–2323. doi:10.1128/JCM.00762-15

24. Lynsdale CL, Santos DJFD, Hayward AD, Mar KU, Htut W, Aung HH, Soe AT, Lummaa V. A standardised faecal collection protocol for intestinal helminth egg counts in Asian elephants, Elephas maximus. Int J Parasitol Parasites Wildl (2015) 4:307–315. doi:10.1016/j.ijppaw.2015.06.001

25. Penzhorn BL. Coccidian oocyst and nematode egg counts of free-ranging African buffalo (Syncerus caffer) in the Kruger National Park, South Africa. J S Afr Vet Assoc (2000) 71:106–108.

26. Herfarth HH, Long MD, Rubinas TC, Sandridge M, Miller MB. Evaluation of a non-invasive method to detect cytomegalovirus (CMV)-DNA in stool samples of patients with inflammatory bowel disease (IBD): a pilot study. Dig Dis Sci (2010) 55:1053–1058. doi:10.1007/s10620-010-1146-0

27. Gillespie TR, Nunn CL, Leendertz FH. Integrative approaches to the study of primate infectious disease: implications for biodiversity conservation and global health. Am J Phys Anthropol (2008) Suppl 47:53–69. doi:10.1002/ajpa.20949

28. Hepworth MR, Daniłowicz-Luebert E, Rausch S, Metz M, Klotz C, Maurer M, Hartmann S. Mast cells orchestrate type 2 immunity to helminths through regulation of tissue-derived cytokines. Proc Natl Acad Sci USA (2012) 109:6644–6649. doi:10.1073/pnas.1112268109

29. Anthony RM, Rutitzky LI, Urban JF, Stadecker MJ, Gause WC. Protective immune mechanisms in helminth infection. Nat Rev Immunol (2007) 7:975–987. doi:10.1038/nri2199

30. Bullen TF, Forrest S, Campbell F, Dodson AR, Hershman MJ, Pritchard DM, Turner JR, Montrose MH, Watson AJM. Characterization of epithelial cell shedding from human small intestine. Lab Invest (2006) 86:1052–1063. doi:10.1038/labinvest.3700464

31. Swamy M, Jamora C, Havran W, Hayday A. Epithelial decision makers: in search of the “epimmunome.” Nat Immunol (2010) 11:656–665. doi:10.1038/ni.1905

32. Knoop KA, Newberry RD. Goblet cells: multifaceted players in immunity at mucosal surfaces. Mucosal Immunol (2018) doi:10.1038/s41385-018-0039-y

33. Crawley JAH, Chapman SN, Lummaa V, Lynsdale CL. Testing storage methods of faecal samples for subsequent measurement of helminth egg numbers in the domestic horse. Vet Parasitol (2016) 221:130–133. doi:10.1016/j.vetpar.2016.03.012

34. Su C, Cao Y, Kaplan J, Zhang M, Li W, Conroy M, Walker WA, Shi HN. Duodenal Helminth Infection Alters Barrier Function of the Colonic Epithelium via Adaptive Immune Activation ∇. Infect Immun (2011) 79:2285–2294. doi:10.1128/IAI.01123-10

35. Midha A, Schlosser J, Hartmann S. Reciprocal Interactions between Nematodes and Their Microbial Environments. Front Cell Infect Microbiol (2017) 7: doi:10.3389/fcimb.2017.00144

