## Supplementary figures and images for "A novel non-invasive method to detect gut barrier related changes during a gastrointestinal nematode infection"

### Supplementary file 1

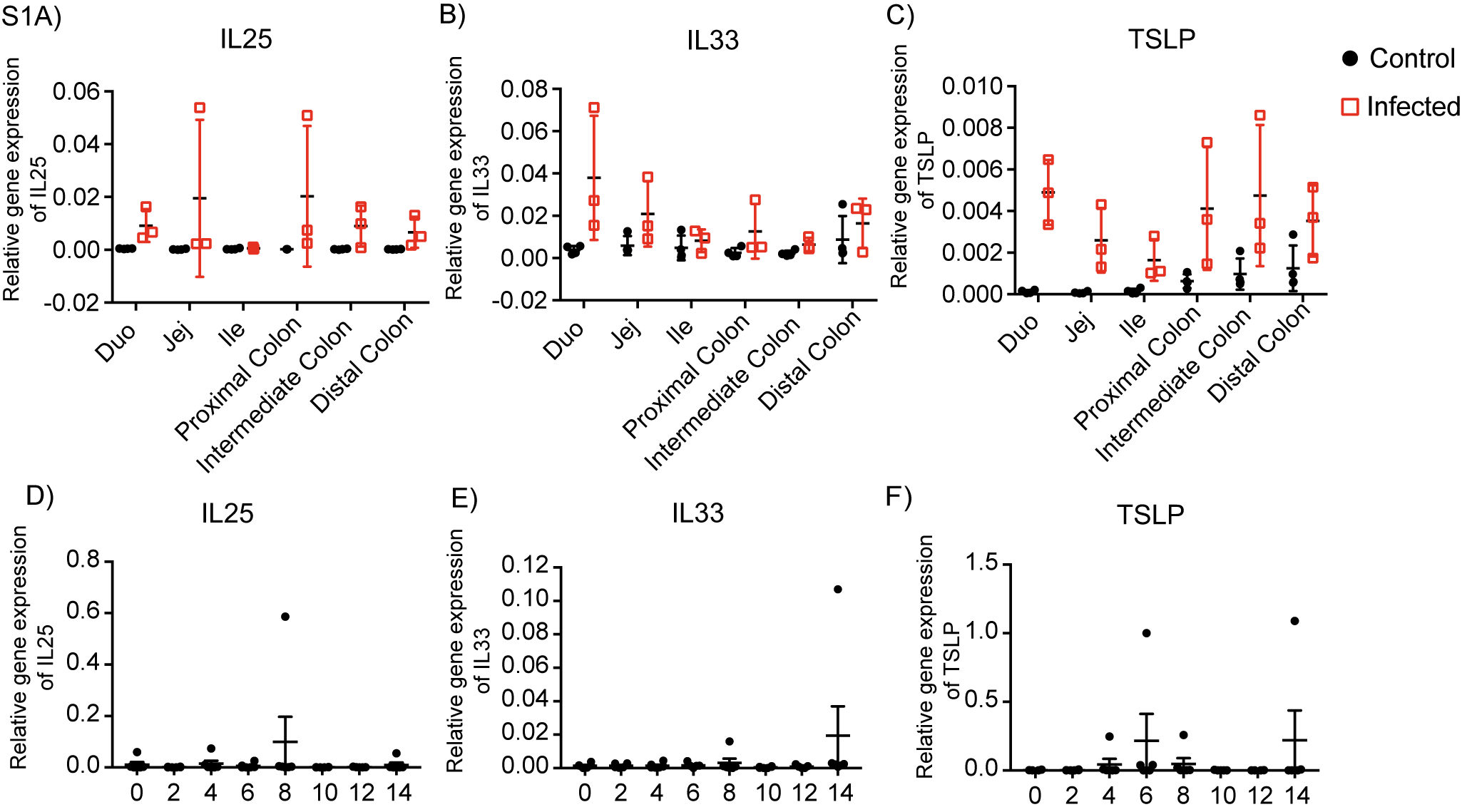
