## Supplementary material for "A novel non-invasive method to detect gut barrier related changes during a gastrointestinal nematode infection"

|  | **Exfoliated cells** | |  | **Homogenized stool** | |
| --- | --- | --- | --- | --- | --- |
| **Gene** | **10ng cDNA (Ct)** | **100ng cDNA (Ct)** |  | **10ng cDNA (Ct)** | **100ng cDNA (Ct)** |
| βactin | 19-25 | 18-21 |  | 27-31 | 22-27 |
