## Supplementary material for "A novel non-invasive method to detect gut barrier related changes during a gastrointestinal nematode infection"

|  | **Exfoliated cells** | | |
| --- | --- | --- | --- |
| **Gene** | **10ng cDNA (Ct)** |  | **100ng cDNA (Ct)** |
| βactin | 23-24 |  | 19 |
| GUSb | 38 |  | 25-29 |
| GAPDH | 32-34 |  | 27-28 |
| HPRT | 30 |  | 26 |
