## Supplementary material for "A novel non-invasive method to detect gut barrier related changes during a gastrointestinal nematode infection"

| Mouse ID | GPS  latitude | GPS  longitude | Sex | Species | Worms | *Hymenolepis*  *microstoma* | *Catenotaenia*  *pusilla* | *Taenia martis* | *Mastophorus muris* | | *Trichurismuris* | *Heterakis* | *Syphacia obvelata* | | *Aspiculuris tetraptera* |
| --- | --- | --- | --- | --- | --- | --- | --- | --- | --- | --- | --- | --- | --- | --- | --- |
| AA_0268 | 52,42473 | 13,70713 | M | Mus musculus | TRUE | 0 | 0 | 0 | | 2 | 9 | 0 | 12 | 0 | |
| AA_0270 | 52,42473 | 13,70713 | F | Mus musculus | TRUE | 0 | 2 | 0 | | 0 | 9 | 0 | 0 | 85 | |
| AA_0290 | 52,26053 | 13,8756 | F | Mus musculus | FALSE | 0 | 0 | 0 | | 0 | 0 | 0 | 0 | 0 | |
| AA_0299 | 52,38064 | 14,095413 | M | Mus musculus | TRUE | 0 | 0 | 0 | | 0 | 83 | 24 | 0 | 5 | |
| AA_0300 | 52,91376 | 14,116181 | F | Mus musculus | TRUE | 0 | 0 | 0 | | 0 | 86 | 17 | 0 | 0 | |
| AA_0318 | 52,91376 | 14,116181 | F | Mus musculus | TRUE | 0 | 0 | 0 | | 0 | 15 | 19 | 76 | 0 | |
| AA_0325 | 53,49615 | 13,440793 | M | Mus musculus | TRUE | 3 | 0 | 0 | | 0 | 0 | 0 | 7 | 0 | |
| AA_0337 | 52,27667 | 14,0817 | F | Mus musculus | TRUE | 0 | 0 | 0 | | 0 | 4 | 0 | 115 | 66 | |
| AA_0343 | 51,60206 | 14,01867 | F | Mus musculus | FALSE | 0 | 0 | 0 | | 0 | 0 | 0 | 0 | 0 | |
| AA_0344 | 53,49615 | 13,440793 | F | Mus musculus | TRUE | 1 | 0 | 3 | | 0 | 3 | 1 | 0 | 0 | |
| AA_0354 | 53,49615 | 13,440793 | F | Mus musculus | FALSE | 0 | 0 | 0 | | 0 | 0 | 0 | 0 | 0 | |
| AA_0355 | 53,49615 | 13,440793 | F | Mus musculus | TRUE | 1 | 0 | 0 | | 17 | 4 | 0 | 0 | 7 | |
| AA_0371 | 52,71519 | 13,957859 | F | Mus musculus | FALSE | 0 | 0 | 0 | | 0 | 0 | 0 | 0 | 0 | |
| AA_0398 | 53,27386 | 13,486372 | F | Mus musculus | FALSE | 0 | 0 | 0 | | 0 | 0 | 0 | 0 | 0 | |
| AA_0409 | 52,74772 | 14,180929 | F | Mus musculus | FALSE | 0 | 0 | 0 | | 0 | 0 | 0 | 0 | 0 | |
| AA_0439 | 52,73734 | 14,075887 | M | Mus musculus | FALSE | 0 | 0 | 0 | | 0 | 0 | 0 | 0 | 0 | |
| AA_0445 | 52,75081 | 14,227153 | F | Mus musculus | FALSE | 0 | 0 | 0 | | 0 | 0 | 0 | 0 | 0 | |
| AA_0450 | 53,32108 | 13,6506 | F | Mus musculus | FALSE | 0 | 0 | 0 | | 0 | 0 | 0 | 0 | 0 | |
| AA_0489 | 53,84506 | 11,4812 | F | Mus musculus | FALSE | 0 | 0 | 0 | | 0 | 0 | 0 | 0 | 0 | |
| AA_0490 | 53,84506 | 11,4812 | M | Mus musculus | FALSE | 0 | 0 | 0 | | 0 | 0 | 0 | 0 | 0 | |
| AA_0497 | 53,94049 | 11,6069 | M | Mus musculus | TRUE | 0 | 0 | 0 | | 0 | 0 | 10 | 0 | 284 | |
| AA_0500 | 53,94049 | 11,6069 | M | Mus musculus | TRUE | 0 | 0 | 0 | | 0 | 0 | 0 | 86 | 118 | |
| AA_0505 | 53,67821 | 12,59032 | F | Mus musculus | TRUE | 0 | 0 | 0 | | 0 | 52 | 0 | 0 | 0 | |
| AA_0510 | 53,36395 | 12,38819 | F | Mus musculus | FALSE | 0 | 0 | 0 | | 0 | 0 | 0 | 0 | 0 | |
